# *Ctcf* Haploinsufficiency Mediates Intron Retention in A Tissue-specific Manner

**DOI:** 10.1101/851923

**Authors:** Adel B Alharbi, Ulf Schmitz, Amy D Marshall, Darya Vanichkina, Rajini Nagarajah, Melissa Vellozzi, Justin JL Wong, Charles G Bailey, John EJ Rasko

## Abstract

CTCF is a master regulator of gene transcription and chromatin organization with occupancy at thousands of DNA target sites. CTCF is essential for embryonic development and somatic cell viability and has been characterized as a haploinsufficient tumor suppressor. Increasing evidence demonstrates CTCF as a key player in several alternative splicing (AS) regulatory mechanisms, including transcription elongation, regulation of splicing factors, and epigenetic regulation. However, the genome-wide impact of *Ctcf* dosage on AS has not been investigated. We examined the effect of *Ctcf* haploinsufficiency on gene expression and AS in multiple tissues from *Ctcf* hemizygous (*Ctcf*^+/-^) mice. Distinct tissue-specific differences in gene expression and AS were observed in *Ctcf*^+/-^ mice compared to wildtype mice. We observed a surprisingly large number of increased intron retention (IR) events in *Ctcf*^+/-^ liver and kidney, specifically in genes associated with cytoskeletal organization, splicing and metabolism. This study provides further evidence for *Ctcf* dose-dependent and tissue-specific regulation of gene expression and AS. Our data provide a strong foundation for elucidating the mechanistic role of CTCF in AS regulation and its biological consequences.

## Introduction

CCCTC-binding factor (CTCF) is a highly conserved, multivalent, 11-zinc finger DNA- and RNA-binding protein, which occupies thousands of conserved target sites distributed across vertebrate genomes and co-ordinates topologically associating domain (TAD) formation [1–5]. Numerous studies have shown that CTCF regulates diverse biological functions including transcriptional activation and repression, insulation, high-order chromatin organization, X-chromosome inactivation, pre-mRNA splicing, and DNA methylation [6, 7]. CTCF binding occurs in a tissue-specific manner, and the binding sites themselves are highly conserved among different species [2–5,8,9]. Over 30% and 50% of all CTCF binding sites are located in intronic and intergenic regions, respectively [2,5,9].

The ubiquitously expressed transcription factor CTCF exhibits differential expression in mammalian tissues [8,10,11], which require specific CTCF levels for growth, differentiation and development [12–14]. Homozygous deletion of *Ctcf* causes early embryonic lethality in mice [15], while mice harboring a hemizygous deletion of *Ctcf* (*Ctcf*^+/-^) exhibit delayed post-natal growth and development [15, 16]. Sustained *Ctcf* haploinsufficiency in mice has various pathophysiological implications including spontaneous widespread tumor formation in diverse tissues as well as genome-wide aberrant hypermethylation [17]. Consistent with that, we have shown that CTCF acts as a haploinsufficient tumor suppressor [16,18,19]. Nevertheless, the underlying link between *Ctcf* haploinsufficiency and spontaneous tumor formation remains to be fully elucidated.

Compelling evidence has linked CTCF to alternative splicing (AS) regulation due to its direct or indirect role in modulating splicing decisions via complex mechanisms including RNA Polymerase II (Pol II) elongation, transcriptional regulation of splicing factors, DNA methylation and chromatin organization [20–25]. Despite its potential importance, the impact of altered CTCF dosage on AS has not been investigated at a global scale. More than 90% of human multi-exonic genes undergo AS in a tissue-specific manner [26, 27]. While normal AS contributes to transcriptome and proteome diversity, aberrant AS can have deleterious effects leading to the development of several pathological complications including cancer [28–30]. Therefore, identifying potential modulators of AS is critical to fully comprehend the etiology and molecular pathophysiology of cancer.

In this study, we conducted transcriptomic analyses in *Ctcf*^+/-^ mice to explore the impact of *Ctcf* haploinsufficiency in AS regulation in spleen, brain, kidney, muscle and liver. We found that *Ctcf* haploinsufficiency induces changes to gene expression and AS that are strikingly tissue-specific. Intron retention (IR), for example, is highly differentially regulated in *Ctcf* haploinsufficient liver and kidney compared to wildtype (WT) mice. Differential IR could be mediated by Ctcf binding sites located up- and downstream of retained introns.

## Results

### Ctcf mediates tissue-specific gene expression and alternative splicing

Transcriptomic and proteomic studies have revealed that CTCF is expressed in all mammalian tissues at levels that vary by up to 20-fold [10, 11]. To assess the dosage-dependent impact of CTCF on gene regulation and AS we examined a *Ctcf* haploinsufficient mouse model harboring a hemizygous deletion of *Ctcf*. In this model, the entire coding region of one *Ctcf* allele is replaced with an expression cassette containing the *pgk* promoter and neomycin gene (*Ctcf*^+/pgkneo^, herein referred to as *Ctcf*^+/-^ for simplicity) (Figure 1A, Supplementary Figure 1). We selected five tissues from different body systems for which the relative CTCF expression between tissues is consistent for human and mouse (Supplementary Figure 2), including lymphatic (spleen), nervous (brain), urinary (kidney), muscular (quadriceps femoris) and digestive (liver) systems. We isolated all five tissues from 11-week-old female mice comprising *Ctcf*^+/-^ and WT littermates, and validated the reduction of *Ctcf* mRNA (by 36-41%) and Ctcf protein (the 130 kDa species by 18-59%) expression in *Ctcf*^+/-^ mice by RT-qPCR and Western blotting, respectively (Supplementary Figure 3). Interestingly, we observed *Ctcf* expression compensation at the protein level in all tissues except spleen, in which the Ctcf concentration was less than 40% of WT Ctcf protein levels. Apart from the spleen, the increase in Ctcf expression from DNA to RNA and protein suggests that a post-transcriptional dosage compensation of *Ctcf* may have occurred (Figure 1B).

**Figure 1:**
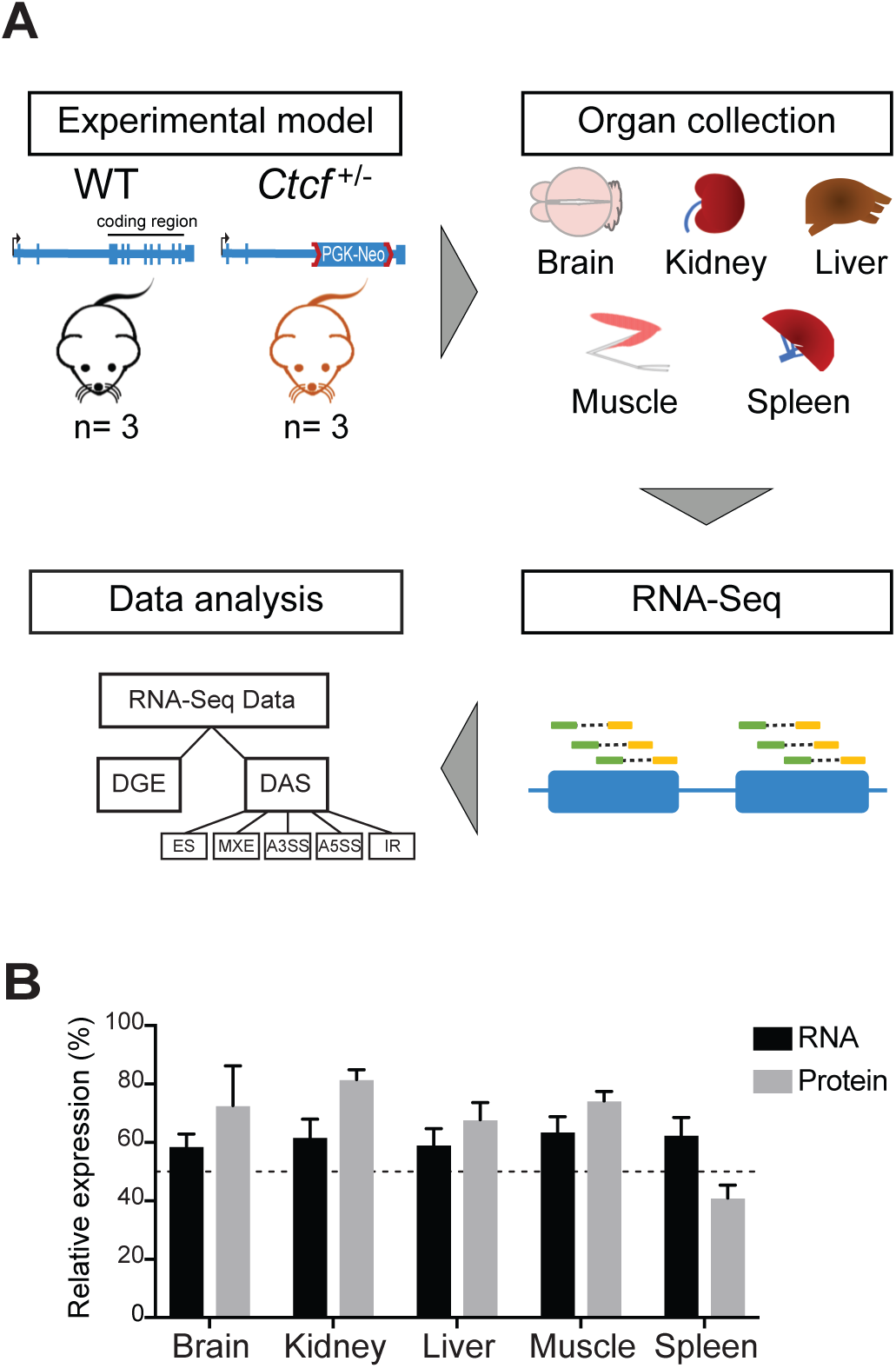
*Ctcf* haploinsufficiency as a model to study Ctcf-mediated transcriptome changes. **(A)** Schematic illustration of the study design highlighting the mouse model, selected organs and types of data analyses. The RNA-seq data analyses performed in this study were differential gene expression (DGE) and differential alternative splicing (DAS) including exon sipping (ES), mutually exclusive exon (MXE), alternative 3′ splice site (A3SS), alternative 5′ splice site (A5SS) and intron retention (IR). **(B)** Summary of Ctcf mRNA and protein expression in all five tissues from WT and *Ctcf*^+/-^ mice (see Supplementary Figure 3 for specific data and statistical analysis). The bars represent the mean +/- SEM. The dashed line represents the expected level of Ctcf in the *Ctcf* haploinsufficient mice (50%).

We performed transcriptome analysis in all five tissues to examine differential gene regulation and differential AS between WT and *Ctcf*^+/-^ mice. *Ctcf* transcript read counts assessed by RNA-seq in *Ctcf*^+/-^ mice were significantly decreased in all tissues (33-39%) compared to WT mice (Figure 2A). Differential gene expression analysis revealed that *Ctcf* haploinsufficiency exerts distinct tissue-specific effects. We detected at least 400 differentially expressed genes (DEGs) per tissue (a total of 3,746 DEGs in all tissues) (Supplementary Table 1). In all tissues other than liver we detected an overall increase in gene expression (Figure 2B), supporting Ctcf’s role as a transcriptional repressor [6].

**Figure 2:**
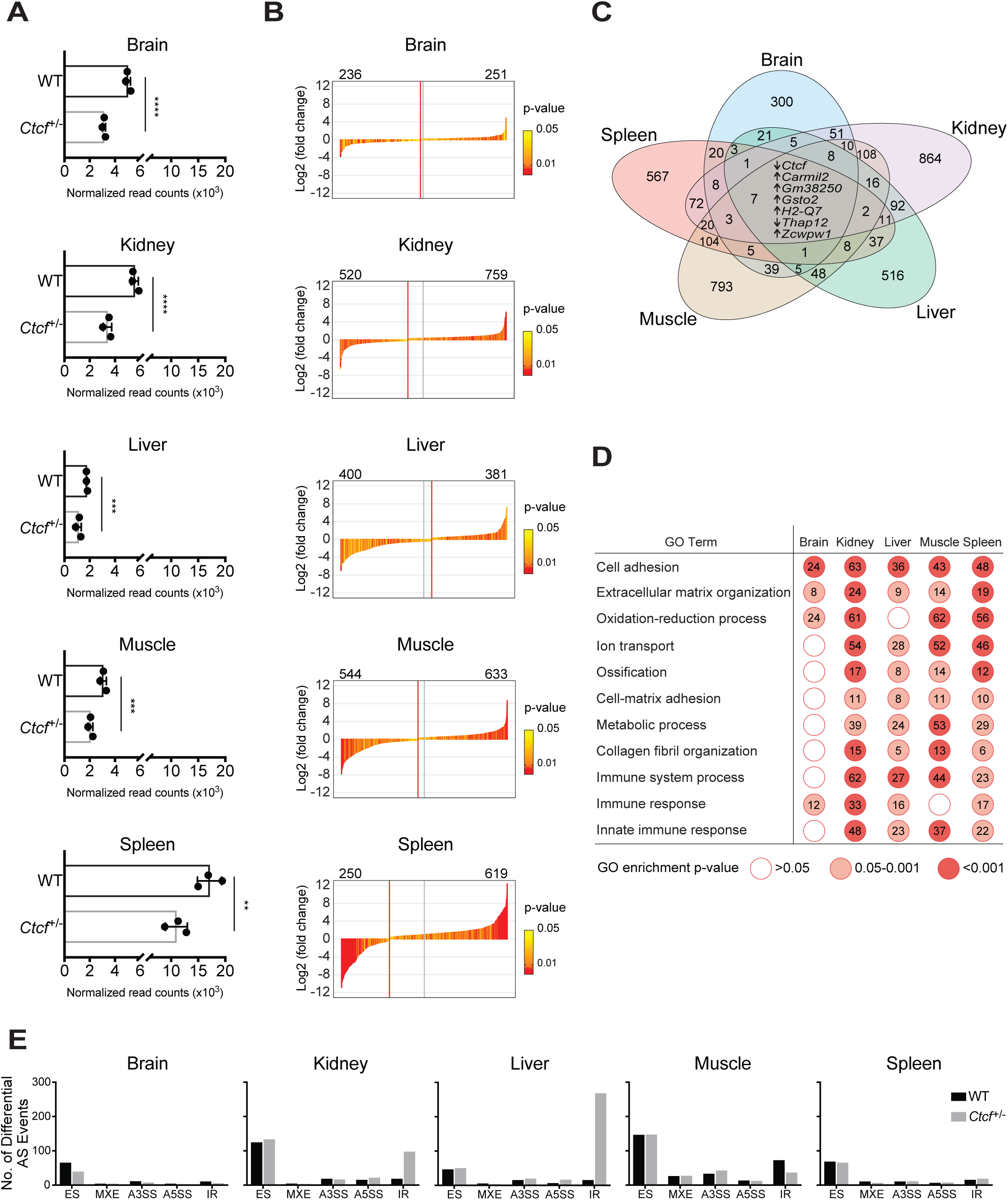
Differential gene expression and alternative splicing in *Ctcf*^+/-^ mice. **(A)** *Ctcf* mRNA expression from five tissues as normalized read counts. The bars represent the mean +/- SEM. Wald test using DESeq2 used to determine significance denoted by ** (*p* < 0.01), *** (p < 0.001), **** (*p* < 0.0001). **(B)** Waterfall plots showing the log2 (fold changes) of all significant DEGs (*p* < 0.05, Wald test using DESeq2) in each tissue. Each vertical bar represents a single DEG which is colored based on the *p*-value. The total numbers of downregulated (left) and upregulated (right) genes are indicated on top of each plot. The central vertical line is drawn at half of the total number of DEGs while the red vertical line indicates the ‘0’ inflection point. Liver was the only tissue not exhibited a reduction in overall gene expression. **(C)** Venn diagram illustrating the overlap of DEGs among the five tissues. Arrows next to the common genes represent the change in their expression as up- or downregulated in all the five tissues. **(D)** Enriched functional annotations associated with DEGs common in at least four out of five tissues. Number of DEGs associated with each GO term is indicated within each circle. **(E)** Bar charts showing the number of significant differential AS events (FDR <0.05, BH) detected in WT and *Ctcf*^+/-^ tissues. *Ctcf*^+/-^ liver exhibited a striking increase in IR. ES - exon sipping; MXE - mutually exclusive exon; A3SS - alternative 3′ splice site, A5SS - alternative 5′ splice site; IR - intron retention.

Apart from *Ctcf*, only six genes (*Carmil2, Gm38250, Gsto2, H2-Q7, Thap12* and *Zcwpw1*) were consistently differentially expressed in all five tissues (Figure 2C). *Carmil2,* located adjacent to *Ctcf* at 8qD3, was upregulated in *Ctcf*^+/-^ mice, which was also observed in a similar hemizygous *Ctcf*^+/-^ mouse model [31]. *ZCWPW1* in humans is located within a CTCF-mediated TAD and contains a CTCF-bound intronic enhancer. A polymorphism (rs1476679) in this enhancer has been associated with increased disease risk in late onset-Alzheimer’s disease by affecting CTCF binding and chromatin topology [32, 33]. A relationship between CTCF and the other four genes has not been established yet.

Gene Ontology (GO)-based annotation enrichment analysis of all the significant DEGs revealed a strong enrichment of tissue-specific biological processes (Supplementary Figure 4). GO terms related to metabolism, immune response, and cellular component organization were commonly affected by *Ctcf* haploinsufficiency (Figure 2D). These data confirm that a decrease in Ctcf dosage through haploinsufficiency can impact gene-expression in a tissue-specific manner.

To examine the effect of *Ctcf* haploinsufficiency on the AS landscape, we analyzed differential AS events in *Ctcf*^+/-^ using rMATS [34]. We found that exon skipping (ES) is the most abundant form of differential AS detected in all tissues (Supplementary Table 2). Notably, we observed a significant increase in IR events in the liver and kidney of *Ctcf*^+/-^ mice (Figure 2E), which is in contrast to all other forms of AS. However, a decrease in Ctcf-mediated IR events was only detected in *Ctcf*^+/-^ muscle. Overall, our analysis shows that *Ctcf* haploinsufficiency not only affects gene expression but also perturbs the AS landscape in a tissue-specific manner.

### Characterization of intron retention in *Ctcf*^+/-^ liver

To confirm that *Ctcf* haploinsufficiency modulates IR in a tissue-specific manner, we used IRFinder, a software package specifically developed for the detection and quantification of IR [35–37]. We confirmed that IR is increased in *Ctcf*^+/-^ liver and kidney with a total of 60 and 43 upregulated IR events, respectively. Only 1 and 5 IR events were downregulated in *Ctcf*^+/-^ liver and kidney, respectively (Figure 3A, Supplementary Table 3). Moreover, we detected that in some mRNA transcripts (5 in liver and 1 in kidney) multiple introns are retained in the same transcript. There were no IR events common to all tissues. Interestingly, we found that a larger number of splicing factors, including *Srsf1*, *Prpf40b*, *Thoc1*, *Khsrp* and *Zpr1,* exhibited retained introns in *Ctcf*^+/-^ liver than other tissues (Supplementary Figure 5).

**Figure 3:**
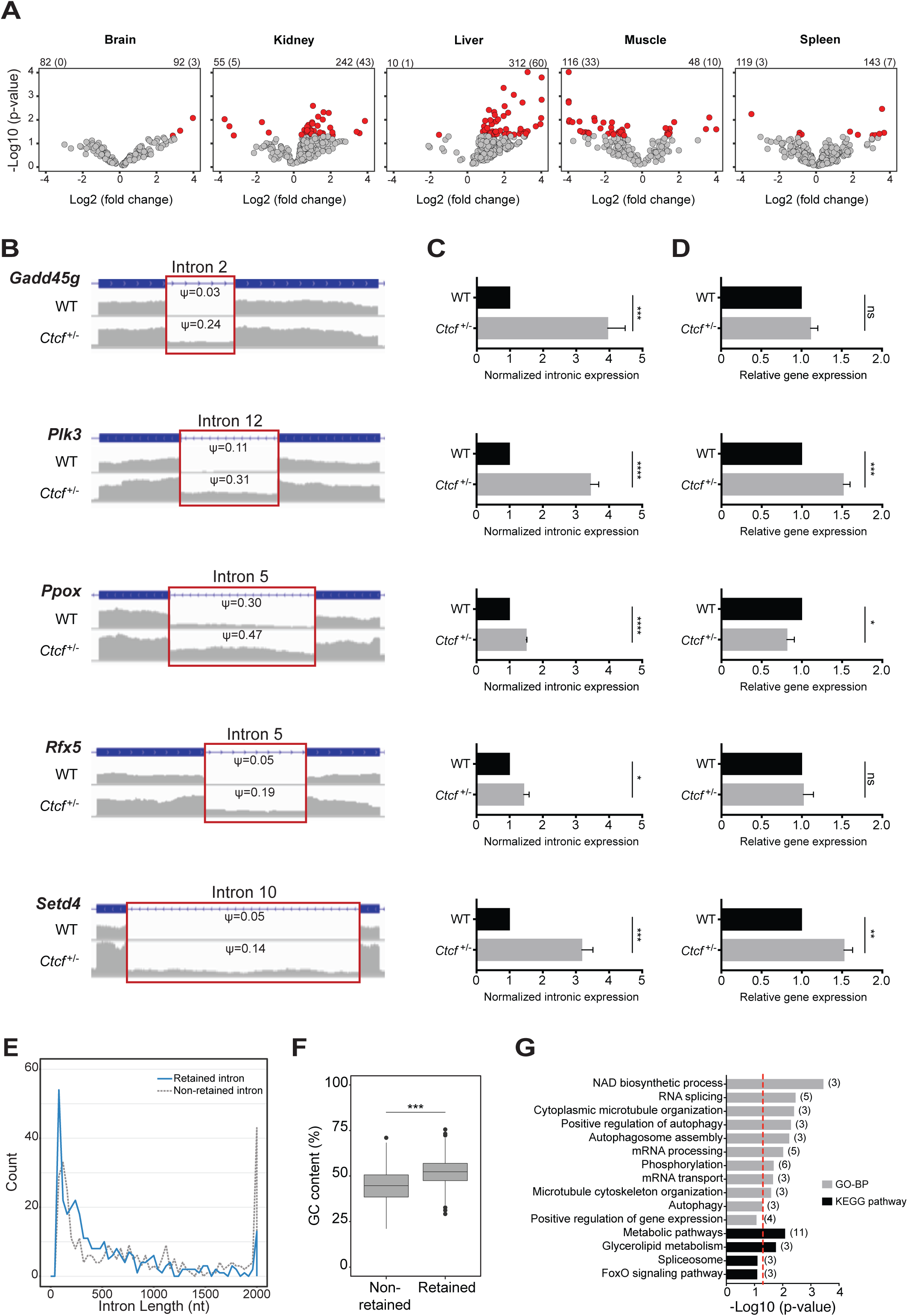
Validation and features of differentially retained introns in liver. **(A)** Volcano plots showing differentially retained introns in mouse tissues. Significant differential IR events (*p* < 0.05, Wald test using IRFinder and DESeq2) are colored in red. The numbers on top represent introns with downregulated (top-left) and upregulated (top-right) IR and the number of significantly differentially retained introns in parentheses. **(B)** Coverage plots of five differentially retained introns selected for validation. The IR-ratio (ψ) of each retained intron is indicated inside the red box. **(C)** Bar charts showing the expression of the same five selected retained introns normalized to the expression of their flanking exons. Unpaired two-tailed Student’s *t*-test was used to determine significance denoted by ns (not significant), * (*p* < 0.05), ** (*p* < 0.01), *** (p < 0.001), **** (*p* < 0.0001). **(D)** Bar charts showing the relative expression of the same five genes harboring the selected differentially retained introns. **(E)** Frequency polygons showing binned length frequencies (binwidth= 40 nt) of differentially retained introns and non-retained introns in *Ctcf*^+/-^ liver. Out of bounds intron length values (>2000bp) were re-scaled and plotted at the maximum value **(F)** Box plot comparing the GC content between retained and non-retained introns in *Ctcf*^+/-^ liver. Significance is denoted by *** (*p* < 0.001, *t*-test). **(G)** Enriched functional annotations associated with genes of the top 60 significantly differentially retained introns in *Ctcf*^+/-^ liver. The dashed vertical red line represents *p*-value=0.05. In parentheses is the number of genes associated with each GO term, where only GO terms with gene number >2 were included.

As *Ctcf* haploinsufficient liver exhibited the most dramatic upregulation of IR, we explored these IR events for further validation and characterization. First, we calculated the IR-ratio for all IR events (Supplementary Figure 6A). The IR-ratio is the proportion of intron-retaining mRNA transcripts compared to all mRNA transcripts from the same gene. Consistent with previous studies, we considered IR events with an IR-ratio ≥ 0.1 as biologically significant [35, 38]. Since we previously showed that IR can reduce the expression of the host gene via nonsense-mediated decay [36], we examined the intronic expression as well as expression of host mRNA. We observed that 65% of differential intron-retaining genes, whether up- or downregulated, exhibited increased gene expression (Supplementary Figure 6B). This suggests that *Ctcf* haploinsufficiency triggers both the upregulation of expression and IR-ratios of intron-retaining genes in liver. We selected five candidates exhibiting increased IR, including *Gadd45g*, *Plk3*, *Rfx5*, *Ppox* and *Setd4* (Figure 3B), for validation by RT-qPCR. We confirmed a significant increase in retained introns in all five genes (Figure 3C), however the impact on host mRNA expression varied (Figure 3D). We have recently summarized all possible fates of intron-retaining transcripts [39, 40]. Apart from their degradation via nonsense-mediated decay, fates include the nuclear detention of intron-retaining transcripts, the synthesis of novel protein isoforms, or their cellular translocation.

We next examined the molecular features of differential IR events identified in *Ctcf*^+/-^ liver and found that retained introns are shorter (Figure 3E) and have a higher GC content than non-retained introns (*p* < 0.0001, Figure 3F), consistent with our previous reports [36,38,41]. To further characterize the liver-specific intron-retaining genes, we performed annotation enrichment analysis and found that the affected genes are associated with biological processes and pathways related to metabolism, cytoskeletal organization, RNA splicing and mRNA processing (Figure 3G). These findings show that *Ctcf* haploinsufficiency induces IR of specific introns in liver genes involved in splicing- and metabolism-related processes.

### Enrichment of Ctcf binding sites proximal to differentially retained introns

CTCF ChIP-seq studies have found that approximately 30% and 50% of CTCF binding sites are located in intronic and intergenic regions, respectively [2,5,9]. It has been shown that the intragenic localization of CTCF binding sites influences pre-mRNA splicing decisions leading to alternative exon splicing [20,21,25]. To determine whether Ctcf binding sites are enriched in the proximity of retained introns in liver, we analyzed a publicly available ChIP-seq data set from C57BL/6 normal mouse liver (E-MTAB-5769) [42].

We examined Ctcf binding sites 200-50,000 nt upstream and downstream of differentially retained introns as well as within each intron. We found that there were more Ctcf binding sites up- and downstream of retained introns compared to expressed non-retained introns (Figure 4A). The fold difference in Ctcf ChIP-seq peaks was higher, both up- and downstream, in regions close to the splice sites (< 2000 nt). In the intron bodies, we observed a non-significant difference in the mean number of Ctcf peaks. The slightly higher mean number of ChIP-seq peaks observed in non-retained introns, could be explained by the fact that retained introns are shorter. These findings suggest that the *Ctcf* haploinsufficiency-induced IR we observed in liver mostly affects short and GC-rich introns which are proximal to Ctcf binding sites.

**Figure 4:**
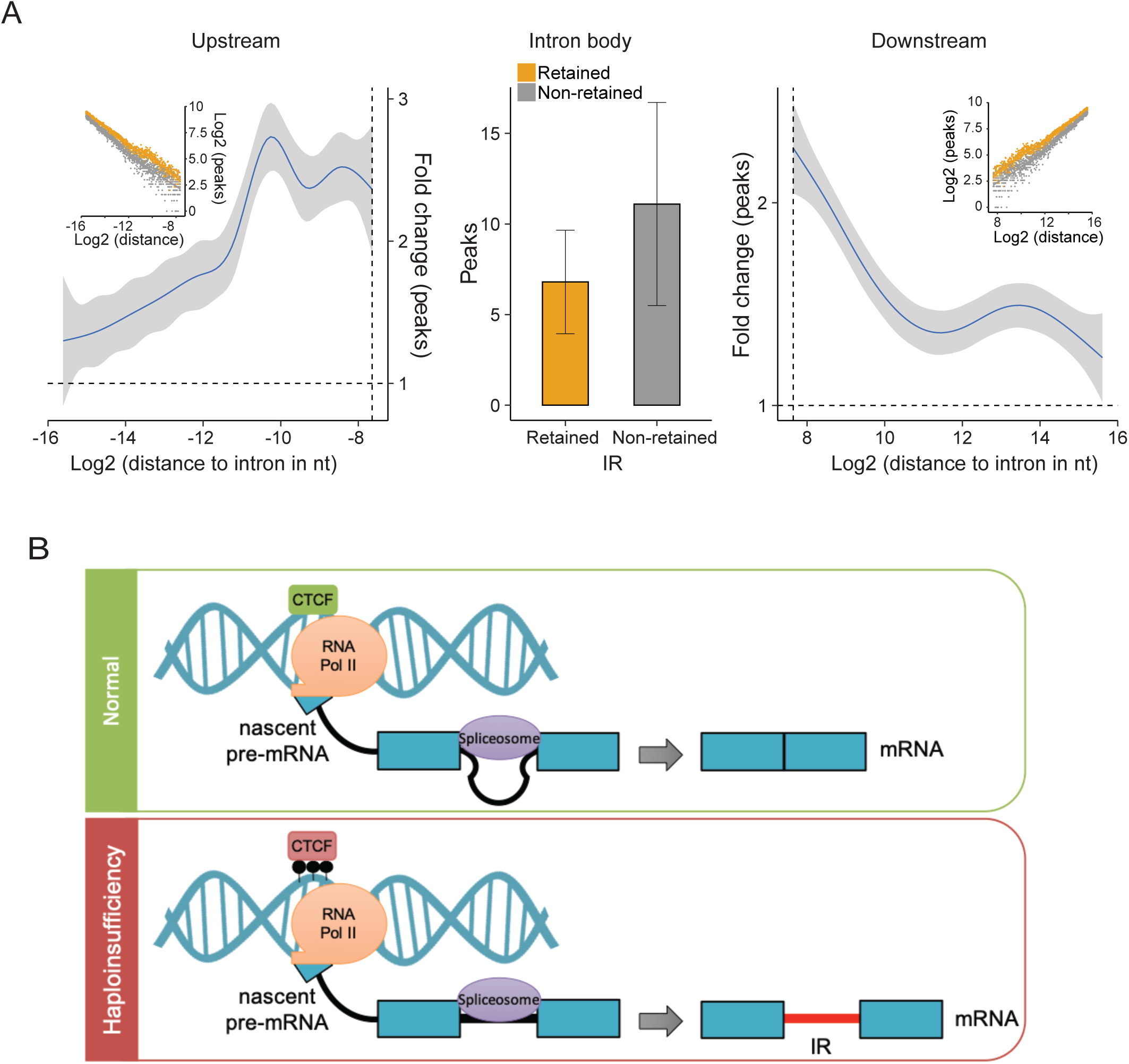
Enrichment of Ctcf binding sites proximal to differentially retained introns. **(A)** Ctcf binding sites upstream (left), downstream (right) and within the body (middle) of retained and non-retained introns. The bar graph in the middle shows the mean number of peaks within the body of introns. The line graphs (left/right) show the distance-dependent fold difference (blue) and standard error bounds (grey) in Ctcf peaks near retained and non-retained introns. The insets show the cumulative number of Ctcf peaks with increasing distance from retained (orange) and non-retained introns (grey). Data points represent 1,000 iterations of 100 randomly selected (non-)retained introns with varying distance to the splice sites. **(B)** Proposed model of *Ctcf* haploinsufficiency-induced IR. In contrast to the normal situation where splicing proceeds, in the *Ctcf* haploinsufficient liver various factors (e.g. RNA Pol II elongation rate, Ctcf expression, Ctcf binding proximal to splice sites and the methylation status of these Ctcf binding sites) contribute to increased IR.

## Discussion

IR is a widespread and conserved mechanism of post-transcriptional gene expression regulation affecting over 80% of all protein coding genes [35]. For instance, we have previously shown that IR regulates differentiation across diverse hematopoietic lineages [36,38,41]. Moreover, the majority of cancer types have abundant IR events that affect their transcriptomes, including genes involved in RNA splicing [43, 44]. Therefore, identifying the potential modulators of AS, particularly IR, is essential to fully comprehend the etiology and molecular pathophysiology of cancer.

Here, we demonstrate that *Ctcf* haploinsufficiency, which has previously been shown to induce spontaneous tumor formation in mice and genome-wide hypermethylation [17], also perturbs the AS landscape. Upon examining a number of tissues, we observed a specific increase of IR in *Ctcf*^+/-^ liver and kidney. Moreover, we observed a similar tissue-specific effect of *Ctcf* haploinsufficiency on gene expression. Given that CTCF expression and DNA occupancy are variable between tissues [2–5,8,9], these differences may govern how CTCF specifically regulates the transcriptome and AS in tissues.

While the impact of CTCF haploinsufficiency on the global AS landscape has not been previously investigated, we utilized this model to examine *in vivo* the global effect of *Ctcf* dose-dependent regulation on AS in specific tissues. Although we used *Ctcf* hemizygous mice, we and others have shown that any sustained decrease in CTCF expression is partially compensated at the protein levels [17, 31]. However, with a less than 50% reduction (except spleen) of Ctcf mRNA and protein expression, we observed tissue-specific differences in gene expression and AS. These data suggest that more prominent effects might be observed with loss-of-function mutations or the inactivation of CTCF, which are commonly observed in cancer [7, 45], or at an even lower CTCF dosage. However, we have previously shown that the enforced genetic ablation of CTCF has a negative impact on somatic cell viability [16].

Although the exact mechanism underlying *Ctcf* haploinsufficiency-mediated IR regulation remains to be characterized, several possibilities can be proposed based on our data and previous studies. An optimal RNA Pol II elongation rate is essential for constitutive splicing [46]; therefore, both slow and fast RNA Pol II elongation rates were observed to promote IR [47]. Slowing RNA Pol II elongation mediated by CTCF promoted exon inclusion in B-lymphoma cell lines, providing the first evidence for a role of CTCF in AS. Binding of CTCF downstream of *CD45* exon 5 leads to decelerating RNA Pol II elongation, thus permitting exon 5 inclusion [20]. This mechanism is tightly regulated by cytosine methylation at CTCF binding sites in the DNA by the methylcytosine dioxygenases TET1 and TET2 [21]. In another example, CTCF binding to the *BDNF2a* locus was shown to be essential for its splicing by inhibiting TET1-induced DNA methylation and the subsequent interaction between TET1, MeCP2 and the splicing factor YB1 [22]. In contrast, DNA methylation at CTCF binding sites located upstream or downstream of alternatively spliced exons was associated with inclusion of exons [24]. Knowing that CTCF regulates DNA methylation and preserves methylation-free regions throughout the genome [48–50], we propose that CTCF regulates AS in a locus-dependent and tissue-specific manner.

CTCF has been recognized as a coordinator of chromatin looping and architecture [51–53]. Of immediate relevance is the observation that the RNA Pol II elongation rate can be controlled by chromatin organization and influence AS decisions [54, 55]. A recent study showed that CTCF-mediated chromatin loops formed between promoters and intragenic regions, particularly at upstream sites of alternatively spliced exons, lead to exon inclusion [25]. However, this mechanism still requires further validation. Overall, these studies suggest a role for CTCF as a key modulator of splicing decisions at individual loci via different molecular and epigenetic mechanisms.

Given its global transcriptional regulation and insulation functions [6], altering CTCF expression could perturb various transcriptional signaling pathways and subsequently AS. In this context, reduced CTCF expression was found to be associated with increased exon exclusion, particularly in those exons located 1 kb upstream of CTCF binding sites in BL41 and BJAB B-lymphoma cells [20]. This suggests that reduction of CTCF expression and the distribution of CTCF binding sites can influence pre-mRNA processing decisions. Consistent with that, we showed that enrichment of Ctcf binding sites upstream and downstream of retained introns in liver is associated with increased IR. From these results we speculate that CTCF may regulate IR through modulating the RNA Pol II elongation rate and methylation status at CTCF target binding sites located proximal to retained introns (Figure 4B).

Our data supports previous observations from our group and others that IR mostly affects introns which are short and GC-rich [36,38,56]. During granulocytic differentiation, some intron-retaining transcripts undergo nonsense-mediated decay, triggered by premature termination codons located within the retained introns [36]. This process can lead to an overall reduction of gene expression consequent to IR. Here, we observed a net increase in gene expression in the liver of *Ctcf*^+/-^ mice, suggesting an alternative predominant fate of intron-retaining transcripts.

We observed that there was an enrichment of genes related to RNA splicing and metabolism among the intron-retaining genes. Splicing factors affected by IR particularly in *Ctcf*^+/-^ liver and kidney such as *Srsf1, Esrp2* and *Prpf40b* have been previously reported to be associated with the regulation of AS including IR [35,57,58]. In addition to splicing, genes involved in metabolic processes were affected by IR in *Ctcf* haploinsufficient liver. We previously reported that *Ctcf* haploinsufficient mice exhibit ∼14% reduction in body weight during post-natal development [16]. Given that the liver plays an essential role in organismal metabolism, we propose that Ctcf-regulated IR in liver may affect metabolic genes and pathways that contribute to this body weight phenotype. *Ctcf* is a haploinsufficient tumor suppressor that induces spontaneous tumor formation in various mouse tissues including liver [17]. The liver exhibited the most abundant IR events observed in our study. Given the fact that IR is upregulated in most human cancers including liver cancer [43], a new link between CTCF haploinsufficiency and hepatocellular carcinoma causation via IR may be possible.

In this study, we have further defined the role of Ctcf in AS regulation and found Ctcf dosage-dependent effects on tissue-specific gene expression. Our analysis of Ctcf binding sites around retained introns in normal liver confirmed their relevance especially in concurrent with short and GC-rich introns. To gain a mechanistic understanding of tissue-specific IR in *Ctcf* haploinsufficiency, additional studies are required that focus on Ctcf DNA occupancy and the interplay with the epigenetic regulation of AS. Moreover, causal links between *Ctcf* haploinsufficiency, IR, and increased tumor formation should be further investigated.

## Materials and methods

### Mouse model, genotyping and organ collection

All mouse experiments were conducted in accordance with the New South Wales Animal Research Act 1985, and the Australian code for the care and use of animals for scientific purposes (8th edition 2013). Animal ethics approval for all mouse experiments was obtained from the Sydney Local Health District Animal Welfare Committee (Protocol number 2016/020). C57BL/6J female mice from Australian BioResources, Australia were used as breeding stock. *Ctcf*^+/−^ mice were originally obtained on a mixed C57BL/6:129SvJ background from the Fred Hutchinson Cancer Research Center (Seattle, WA, USA) as a kind gift from Galina Filappova. These mice were previously backcrossed onto C57Bl/6J mice for at least 10 generations. *Ctcf*^+/-^ mice contain a hemizygous deletion of the entire *Ctcf* coding region and substituted with an expression cassette containing the 3-phosphoglycerate kinase promoter and Neomycin gene flanked by two loxP sites (Pgk-Neo) [15, 16]. Genotyping primers used to distinguish WT and *Ctcf*^+/-^ alleles were Ctcf-WT-5’ (CTCACGCCTGAGATGATCC), Ctcf-WT-3’ (CATGCCATCCTACTGGTGTG) and Ctcf-neo-5’ (TGGGCTCTATGGCTT CTGAG). The resultant amplicons represent the WT allele (519 bp) and PGK-Neo-containing allele (348 bp).

Six healthy female C57BL/6 *Ctcf*^+/+^ and *Ctcf*^+/-^ mice (3 each) were utilized. At 11 weeks of age, two litters, one containing 2xWT and 2x*Ctcf*^+/-^ female mice, the other containing one WT and one *Ctcf*^+/-^ female mouse each were euthanized by carbon dioxide asphyxiation as recommended by the institutional animal welfare guidelines. Organs of interest (brain, kidney, liver, muscle and spleen) were collected and snap-frozen in liquid nitrogen, and stored at −80°C.

### Protein extraction and quantitation

To extract protein from mouse organs, a small piece of the organ was minced in ice-cold RIPA protein lysis buffer (50 mM Tris-HCL (pH 8.0), 150 mM NaCl, 0.1% (w/v) sodium dodecyl sulfate (SDS), 0.5% (w/v) sodium deoxycholate, 1% (v/v) NP-40 and 0.02% (w/v) sodium azide in distilled H_2_O with the addition of 1X EDTA-free SIGMAFAST protease cocktail tablet (Sigma). To quantify protein in the tissue lysates, the Micro BCA Protein Assay Kit (Thermo Fisher Scientific) was used according to the manufacturer’s protocol.

### Western blot, antibodies and densitometry

For Western blot analysis, protein extracts were prepared as equal aliquots (10 μg) and boiled in 2X protein loading buffer containing NuPAGE LDS Sample Buffer and 100 mM Dithiothreitol (DTT) for 5 minutes. The samples were then resolved by 4-12% SDS-PAGE using NuPAGE Bis-Tris Protein Gel system (Thermo Fisher Scientific), and transferred onto a PVDF membrane (Merck Millipore) in a semi-dry transfer apparatus. Rabbit polyclonal anti-CTCF (1:2,000; #3418, Cell Signaling) or mouse monoclonal anti-GAPDH (1:5,000 dilution; ab8245; Abcam) was used as a primary antibody, followed by washing and staining with horseradish peroxidase-conjugated donkey anti-rabbit or anti-mouse IgG secondary antibodies (1:5,000 dilution; Merck Millipore), respectively. A Chemidoc Touch (BioRad) was used to visualize the protein bands on these replicate blots, which were subjected to densitometric analysis using ImageJ software.

### RNA isolation, purification and quantitation

To isolate RNA from WT and *Ctcf*^+/-^ tissues of interest, a small piece of tissue (<30 mg weight) was homogenized with TRIzol reagent (Invitrogen) followed by isopropanol precipitation at −80°C. Next, contaminating DNA was eliminated by using the TURBO DNA-free kit (Invitrogen). The RNA quantitation and quality were assessed by NanoDrop 1000 Spectrophotometer (Thermo Fisher Scientific). Only samples with a 260/280 nm absorbance ratio of 1.9 or above were used. The RNA integrity was evaluated using a 2100 Bioanalyzer with the RNA 6000 Nano kit (Agilent) where samples with an RNA integrity number (RIN) below 7 were excluded.

### RT-qPCR

To determine the relative expression level of Ctcf mRNA, cDNA was synthesized using SuperScript III Reverse Transcriptase (Invitrogen) followed by RNaseOUT treatment (Invitrogen). Next, qPCR was performed on the cDNA with SYBR Green (Bio-Rad) and *Ctcf*-specific forward (CCACCTGCCAAGAAGAGAAG) and reverse (CGACCTGAATGATGGCTGTT) primers, and subsequently run on a CFX96 real-time PCR machine (BioRad). Primers detecting *Hprt* were used to normalize gene expression: forward (AGTGTTGGATACAGGCCAGAC) and reverse (CGTGATTCAAATCCCTGAAGT). The fold-change was calculated using the 2^−ΔΔCt^ method. The *Ctcf* mRNA expression levels in *Ctcf*^+/-^ mouse tissues were normalized to WT to obtain the relative mRNA expression.

### RNA sequencing

A total of thirty RNA samples (five tissues collected from three biological replicates of WT and *Ctcf*^+/-^ mice), which met the quality control parameters described above, were sent to Novogene (Beijing, China) for library preparation (strand-specific, poly-A enriched) and RNA-seq (150 bp paired-end reads, over 100 million reads) using the Illumina HiSeq-PE150 platform.

### Bioinformatic analysis

Initial quality control of the RNA-seq raw data was conducted using FastQC (bioinformatics.babraham.ac.uk/projects/fastqc) and MultiQC (multiqc.info) to check for sequencing quality prior to analysis [59]. After passing the quality control check, reads were mapped to the mouse reference genome GRCm38/mm10 using STAR software [60]. To examine differential gene expression in WT and *Ctcf*^+/-^ tissue samples, read counts were determined using featureCounts 1.5.0-p3 [61] and then subjected to differential gene expression analysis using the R Bioconductor package DESeq2 1.25.15 [62]. Genes with low read counts (<5) across all samples were removed. Differential AS was analyzed using rMATS 4.0.1 [34], while further analysis of differential IR was performed using IRFinder 1.2.5 [35]. Specific IR features such as intron length and GC content were calculated using BEDTools v2.26.0 [63]. GO analysis of differentially expressed genes and intron-retaining genes were performed using the DAVID 6.8 online tool [64].

### Validation of retained-intron candidates

Several candidates from the subset of intron-retaining transcripts in *Ctcf*^+/-^ liver, were selected for validation by RT-qPCR according to stringent filtering parameters from the main IRFinder output file. These filtering parameters included mean IR-ratio ≥0.1, mean intron coverage >75%, mean intron depth ≥5 reads, mean splice junction depth ≥20 reads, and *p*-value of IR-ratio fold-change between WT and *Ctcf*^+/-^ samples <0.05. Further details of these parameters are provided in the IRFinder manual (github.com/williamritchie/IRFinder/wiki). Selected differentially retained introns were also visually inspected using the Integrative Genomics Viewer software (software.broadinstitute.org/software/igv/).

Next, the selected candidates were validated by RT-qPCR using a set of three specific primers (Supplementary Table 4). To measure the relative mRNA expression, two primers were used to amplify sequence spanning the exonic region adjacent to the retained intron (Forward) to cross the exon-exon junction flanking the retained intron (Reverse 1). To measure the retained intron in the target spliced isoform, a third primer was designed to anneal within the intronic region of the retained intron (Reverse 2). Finally, the relative mRNA expression was calculated as described above while intron-retaining transcript expression was calculated after normalizing its expression to the expression of the intron’s flanking exons.

### Statistical analysis

The data from this study was analyzed using different statistical tests relevant to the type of experiment or analysis. Wald test and unpaired two-tailed Student’s t-test (as indicated in figures legends) were used to determine statistical significance. P-values <0.05 were considered as statistically significant. All error bars shown in the figures represent standard error of the mean (SEM) from at least three independent experiments. The GraphPad Prism software version 7 as well as R Bioconductor were used to perform the statistical data analyses.

## Data availability

Data have been submitted to GEO under accession number (GSE140532).

## Acknowledgements

This work was supported by the National Health and Medical Research Council (Grant #1128175 and #1129901 to J.E.J.R.). Financial support was also provided by Tour de Cure (Scott Canner Research Fellowship) to C.G.B. and for research grants to C.G.B. and J.E.J.R; Tour de Rocks project support to C.G.B.; Cancer Council NSW project grants (RG11-12, RG14-09) to J.E.J.R. and C.G.B. A.B.A. is supported by a PhD Scholarship from Umm Al-Qura University in Saudi Arabia. U.S. and J.J.L.W. hold Fellowships from the Cancer Institute of New South Wales. The authors acknowledge the Centenary Institute Animal House staff for animal husbandry.

## Author contributions

The project was conceived by J.E.J.R., C.G.B., A.D.M. and U.S.; experiments were planned by A.D.M. and A.B.A.; experiments were performed by A.D.M., A.B.A. and J.J.L.W.; RNA-seq data were analyzed by A.B.A, U.S and D.V; all other data were analyzed and evaluated by A.B.A.; the mice organs were collected by R.N., A.B.A., A.D.M. and M.V.; the manuscript was written by A.B.A., C.G.B., U.S. and J.E.J.R.

## Supplementary Figure legends

**Supplementary Figure 1:**
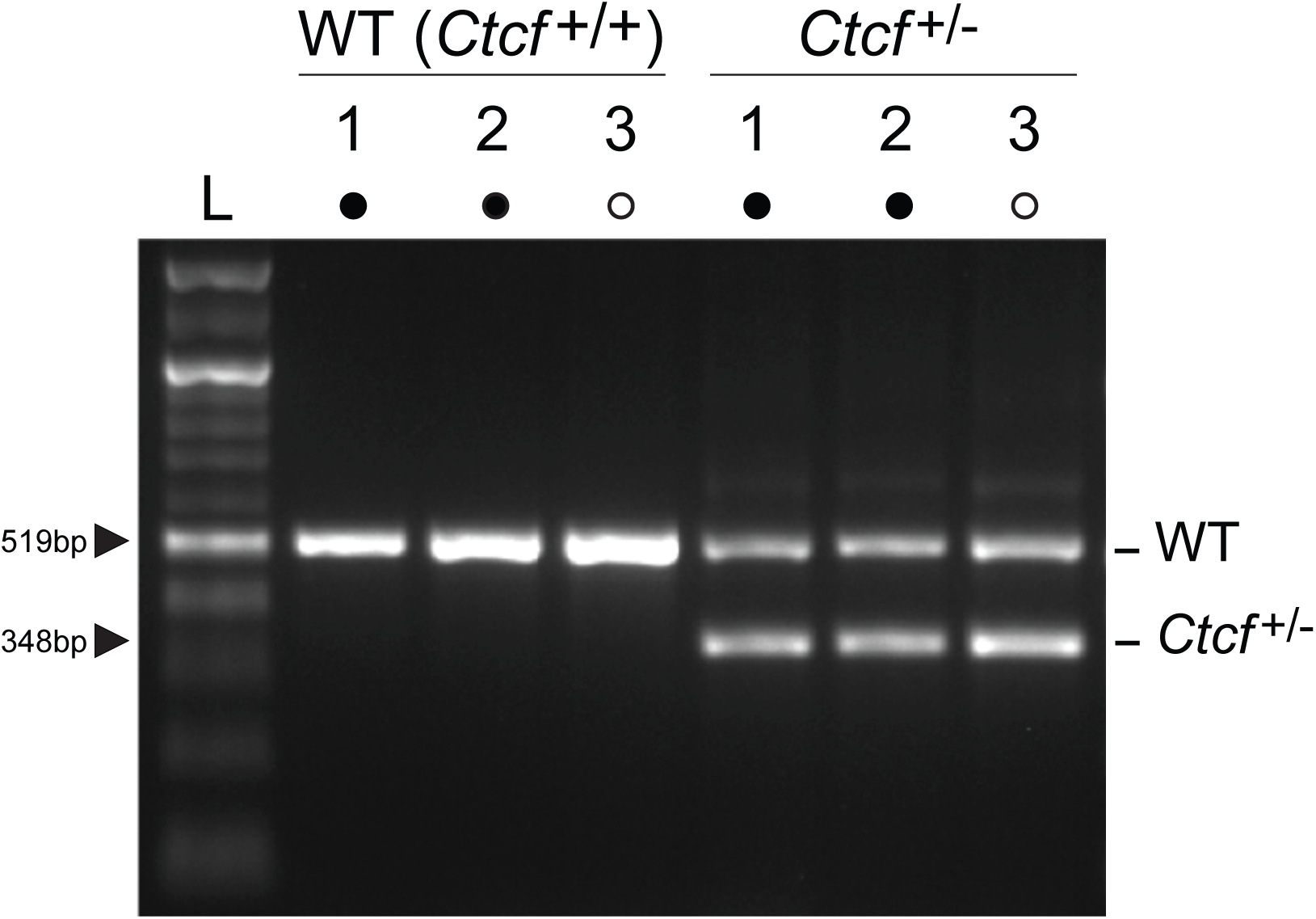
Genotyping of WT (*Ctcf*^+/+^) and *Ctcf*^+/-^ female mice. Using specific primers, WT alleles (519 bp) and *Ctcf*^+/-^ allele containing PGK-Neo (348 bp) were detected. Approximate expected amplicon sizes (in bp) are indicated on the left. L denotes 100bp ladder; filled circles denotes same littermates; open circles denotes same littermates.

**Supplementary Figure 2:**
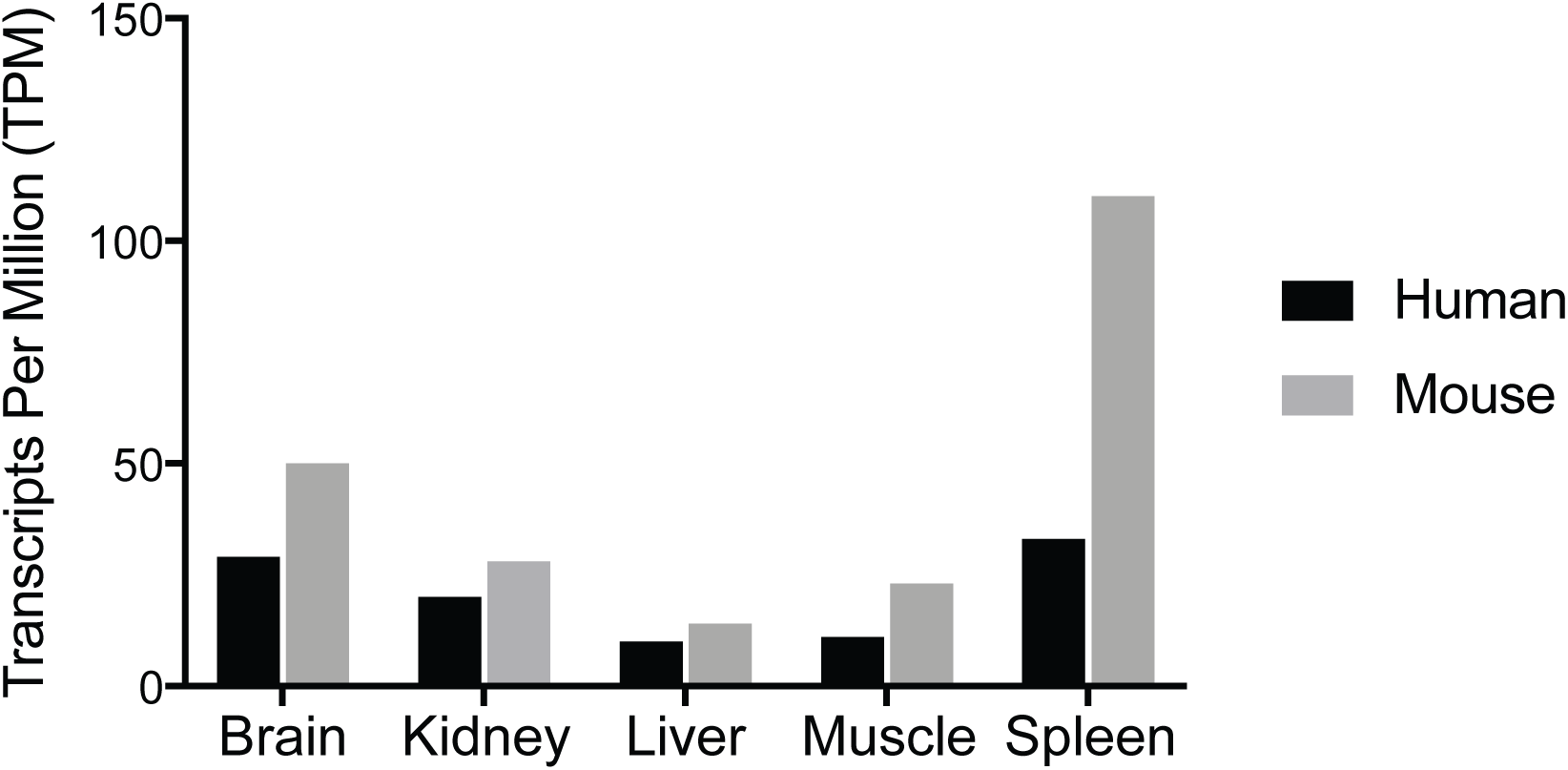
*Ctcf* expression in human and mouse tissues. *Ctcf* expression depicted, as transcripts per million mapped reads, in five human and mouse tissues. The data used to generate this chart were collected from large-scale studies of human and mouse gene expression profiling [10, 11].

**Supplementary Figure 3:**
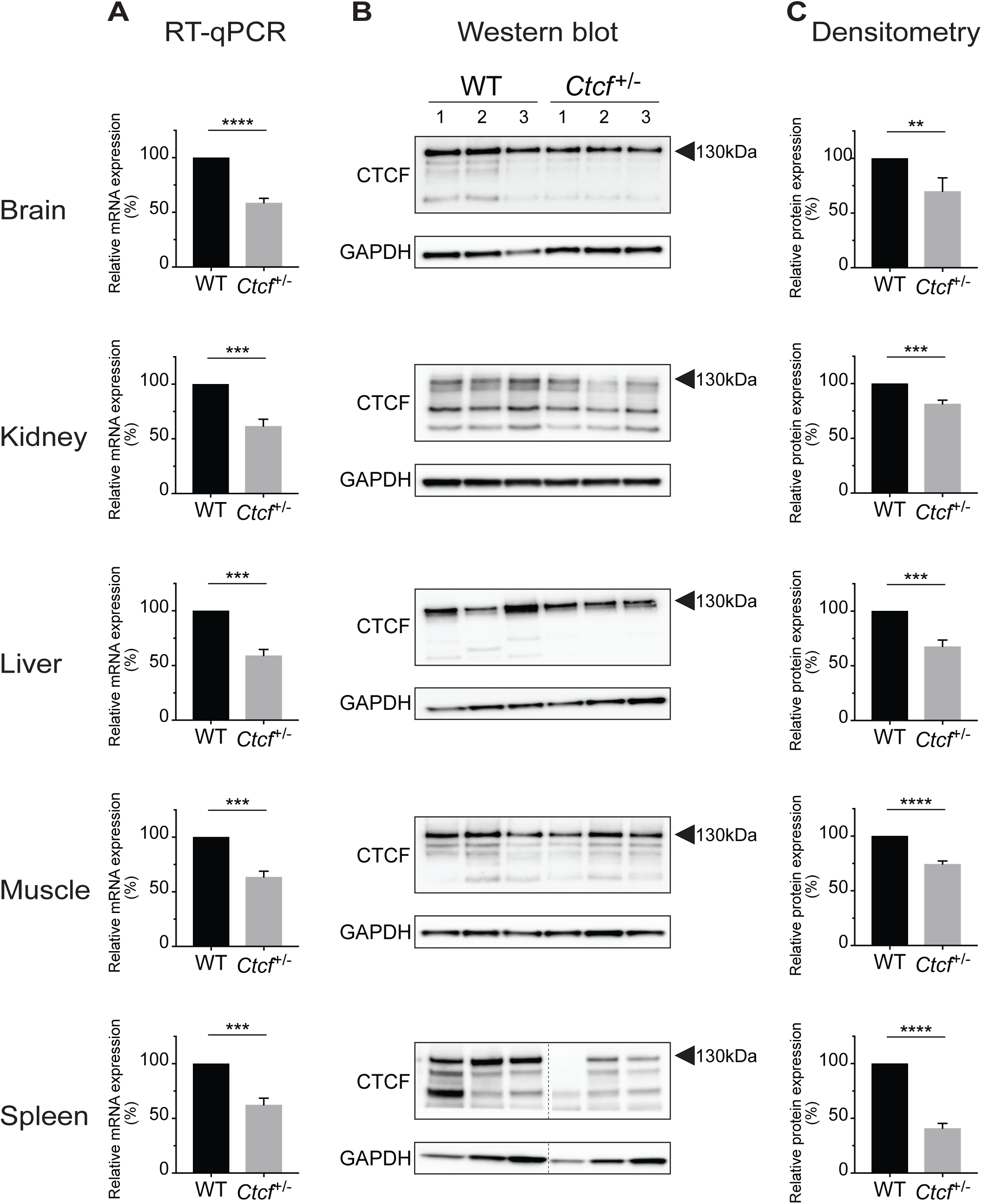
Validation of reduced Ctcf expression in *Ctcf*^+/-^ mice. **(A)** *Ctcf* mRNA and protein **(B)** expression in isolated mouse tissues were assessed by RT-qPCR and Western blotting, respectively. The dashed line in the Western blots from spleen indicates cropping **(C)** Relative Ctcf protein expression was confirmed by ImageJ densitometric analysis of the upper 130 kDa band. *Hprt* mRNA and Gapdh protein were used as loading controls in RT-qPCR and Western blotting, respectively. Statistical significance is denoted by ** (*p* < 0.01), *** (*p* < 0.001), **** (*p* < 0.0001, two-tailed Student’s *t*-test).

**Supplementary Figure 4:**
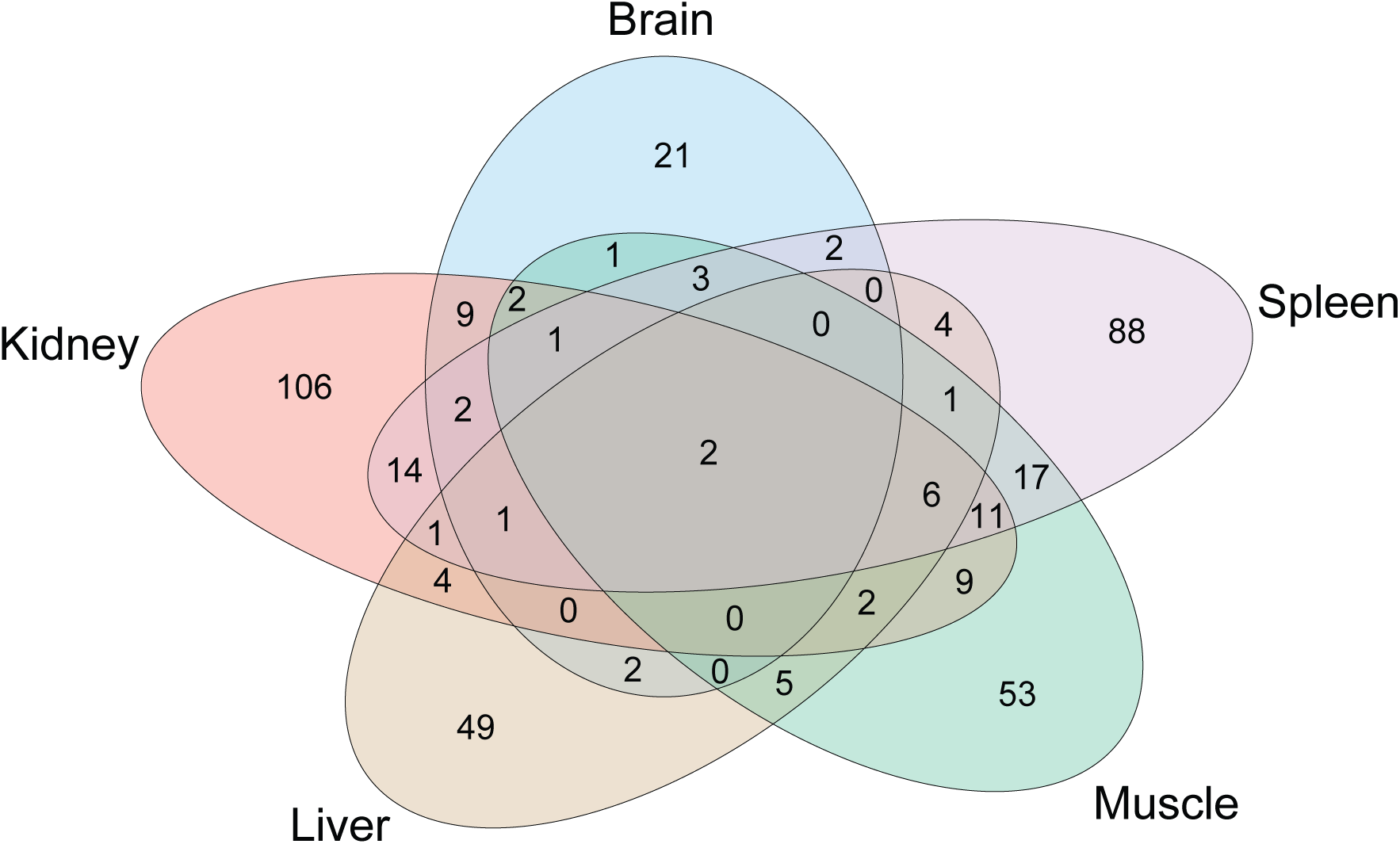
Intersection of significant enriched functional annotations associated with DEGs.

**Supplementary Figure 5:**
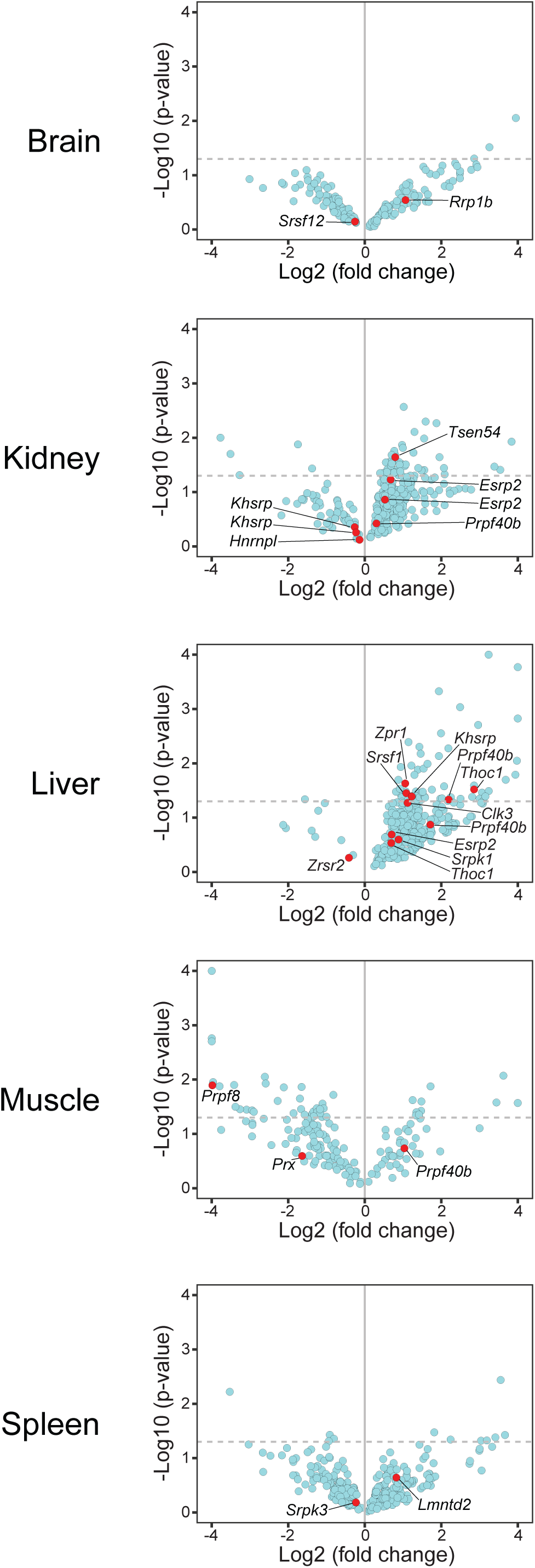
Splicing factors affected by differential IR. Volcano plots showing differentially retained introns in five mouse tissues. RNA splicing factors are highlighted as red dots next to the gene name. Dashed horizontal lines are drawn at *p*-value=0.05 (Wald test using IRFinder and DESeq2).

**Supplementary Figure 6:**
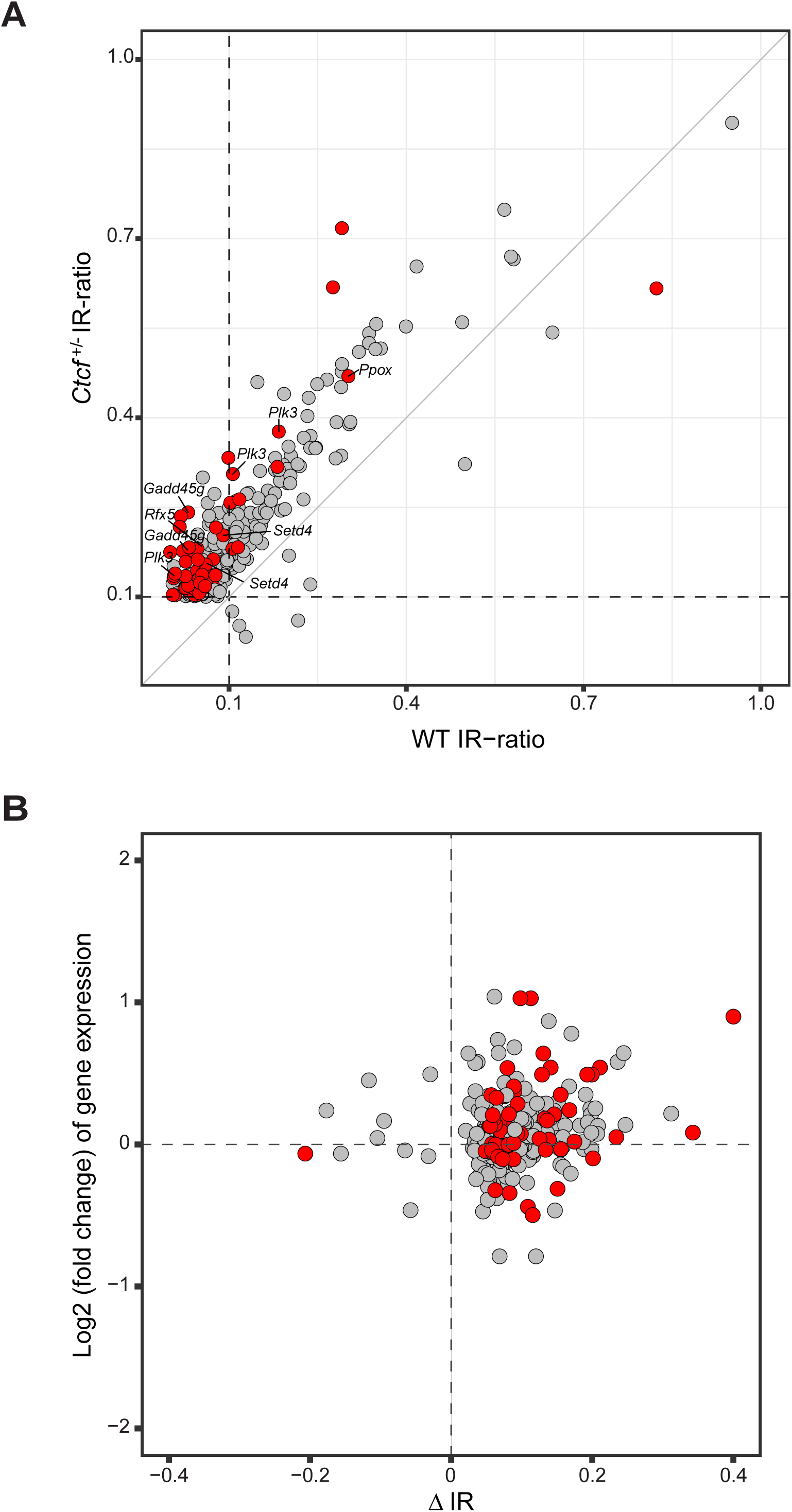
Differential IR and gene expression of intron-retaining genes in *Ctcf^+/-^* liver. **(A)** Scatter plot showing the IR-ratios of differentially retained introns in WT (x-axis) and *Ctcf^+/-^* mice. **(B)** Scatter plot showing changes of the IR-ratio (ΔIR, x-axis) and gene expression (log2 fold change) of intron-retaining genes (y-axis). Significant differentially retained introns (*p* < 0.05, Wald test using IRFinder and DESeq2) are colored in red.

Supplementary Table 1: List of significant DEGs in *Ctcf*^+/-^ mouse tissues.

Supplementary Table 2: List of significant AS events in *Ctcf*^+/-^ mouse tissues.

Supplementary Table 3: List of significant IR events in *Ctcf*^+/-^ liver and kidney.

**Supplementary Table 4:**
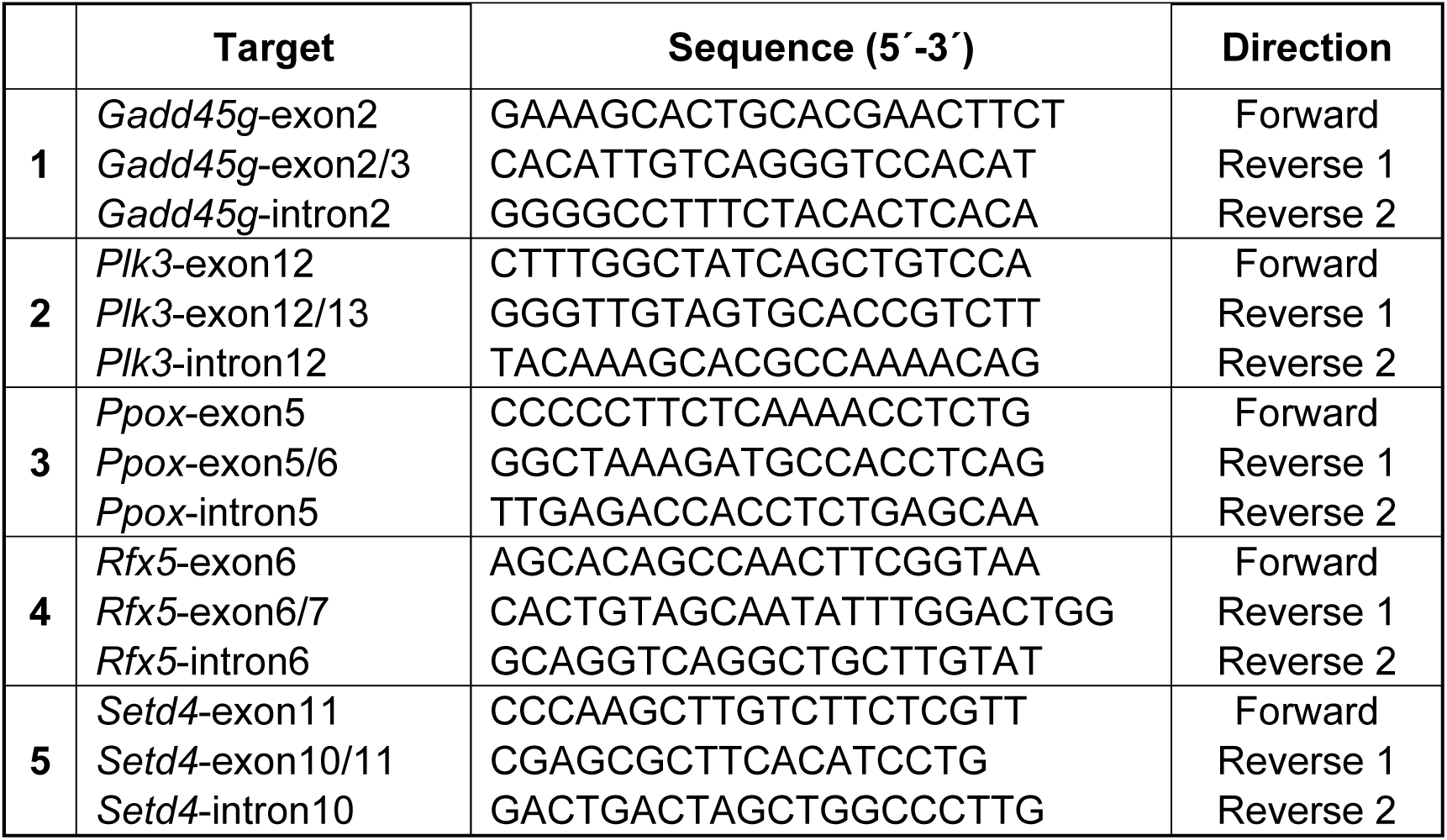
Primers for IR validation by RT-qPCR

